# Functional analysis of African *Xanthomonas oryzae* pv. *oryzae* TALomes reveals a new susceptibility gene in bacterial leaf blight of rice

**DOI:** 10.1101/261313

**Authors:** Tuan Tu Tran, Alvaro L Pérez-Quintero, Issa Wonni, Sara C. D. Carpenter, Yanhua Yu, Li Wang, Jan E. Leach, Valérie Verdier, Sébastien Cunnac, Adam J. Bogdanove, Ralf Koebnik, Mathilde Hutin, Boris Szurek

**Affiliations:** UMR IPME, IRD-CIRAD-Université Montpellier, Montpellier, France; Plant Pathology and Plant-Microbe Biology Section, School of Integrative Plant Science, Cornell University, Ithaca, New York, United States of America; Bioagricultural Sciences and Pest Management, Colorado State University, Fort Collins, United States of America; Agricultural Genetics Institute, Hanoi, Vietnam; Institut de l'Environnement et de Recherches Agricoles, Laboratoire Mixte International, Observatoire des agents phytopathogènes en Afrique de l’Ouest, Bobo Dioulasso, Burkina Faso; State Key Laboratory for Conservation and Utilization of Subtropical Agro-bioresources, The Key Laboratory of Ministry of Education for Microbial and Plant Genetic Engineering, and College of Life Science and Technology, Guangxi University, Nanning, Guangxi, China

## Abstract

Most Xanthomonas species translocate Transcription Activator-Like (TAL) effectors into plant cells where they function like plant transcription factors via a programmable DNA-binding domain. Characterized strains of rice pathogenic *X. oryzae* pv. *oryzae* harbor 9-16 different *tal* effector genes, but the function of only a few of them has been decoded. Using sequencing of entire genomes, we first performed comparative analyses of the complete repertoires of TAL effectors, herein referred to as TALomes, in three *Xoo* strains forming an African genetic lineage different from Asian *Xoo*. A phylogenetic analysis of the three TALomes combined with in silico predictions of TAL effector targets showed that African *Xoo* TALomes are highly conserved, genetically distant from Asian ones, and closely related to TAL effectors from the bacterial leaf streak pathogen *Xanthomonas oryzae* pv. *oryzicola* (*Xoc*). Nine clusters of TAL effectors could be identified among the three TALomes, including three showing higher levels of variation in their repeat variable diresidues (RVDs). Detailed analyses of these groups revealed recombination events as a possible source of variation among TAL effector genes. Next, to address contribution to virulence, nine TAL effector genes from the Malian *Xoo* strain MAI1 and four allelic variants from the Burkinabe *Xoo* strain BAI3, thus representing most of the TAL effector diversity in African *Xoo* strains, were expressed in the TAL effector-deficient *X. oryzae* strain X11-5A for gain-of-function assays. Inoculation of the susceptible rice variety Azucena lead to the discovery of three TAL effectors promoting virulence, including two TAL effectors previously reported to target the susceptibility (*S*) gene *OsSWEET14* and a novel major virulence contributor, TalB. RNA profiling experiments in rice and *in silico* prediction of EBEs were carried out to identify candidate targets of TalB, revealing *OsTFX1*, a bZIP transcription factor previously identified as a bacterial blight *S* gene, and *OsERF#123*, which encodes a subgroup IXc AP2/ERF transcription factor. Use of designer TAL effectors demonstrated that induction of either gene resulted in greater susceptibility to strain X11-5A. The induction of *OsERF#123* by BAI3Δ1, a *talB* knockout derivative of BAI3, carrying these designer TAL effectors increased virulence of BAI3Δ1 validating OsERF#123 as a new, bacterial blight *S* gene.

**Author Summary:** The ability of most *Xanthomonas* plant pathogenic bacteria to infect their hosts relies on the action of a specific family of proteins called TAL effectors, which are transcriptional activators injected into the plant by the bacteria. TAL effectors enter the plant cell nucleus and bind to the promoters of specific plant genes. Genes that when induced can benefit pathogen multiplication or disease development are called susceptibility (*S*) genes. Here, we perform a comparative analysis of the TAL effector repertoires of three strains of *X. oryzae* pv. *oryzae*, which causes bacterial leaf blight of rice, a major yield constraint in this staple crop. Using sequencing of entire genomes, we compared the large repertoires of TAL effectors in three African *Xoo* strains which form a genetic lineage distinct from Asian strains. We assessed the individual contribution to pathogen virulence of 13 TAL effector variants represented in the three strains, and identified one that makes a major contribution. By combining host transcriptome profiling and TAL effector binding sites prediction, we identified two targets of this TAL effector that function as *S* genes, one previously identified, and one, new *S* gene. We validated the new *S* gene by functional characterization using designer TAL effectors. Both *S* genes encode transcription factors and can therefore be considered as susceptibility hubs for pathogen manipulation of the host transcriptome. Our results provide new insights into the diversified strategies underlying the roles of TAL effectors in promoting plant disease.

## Introduction

*X. oryzae* pv. *oryzae* (*Xoo*) and *X. oryzae* pv. *oryzicola* (*Xoc*) respectively cause bacterial leaf blight (BLB) and bacterial leaf streak (BLS) of rice, two important constraints for production worldwide, causing yield losses ranging from 0-30% for BLS and up to 50% for BLB [1]. Although originally discovered in Asia, both diseases were reported in West Africa in the 1980’s. Since then, reports of both BLB and BLS in different countries across the continent have increased, illustrating their potential effect on African rice production [2]. Although closely related genetically, *Xoo* and *Xoc* cause different symptoms in early stages of infection and colonize the host in different ways. While *Xoo* is a vascular pathogen, entering rice leaves via hydathodes or wounds and colonizing the xylem parenchyma [3], *Xoc* is an intercellular pathogen infecting leaves through stomata and multiplying in the mesophyll apoplast. Genetic diversity analysis revealed that *Xoo* can be classified into distinct genetic lineages according to geographical distribution where Asian and African strains define two separate clades. Intriguingly, African *Xoo* strains seem most closely related to and share several features with *Xoc* [4-6]. Pathogenicity assays grouped African *Xoo* into three novel races [5], and other studies indicated that they trigger specific resistance responses [7].

To successfully colonize their host, both *Xoc* and *Xoo* rely to some extent on host transcriptional changes caused by Transcription Activator-Like (TAL) effectors. TAL effectors form a unique class of type III effectors conserved in many *Xanthomonas spp.* [8]. Upon injection via the type III secretion system, TAL effectors localize to the host nucleus where they directly bind to effector-specific DNA sequences and activate transcription at those locations [9]. When induction of targeted genes promotes bacterial colonization *in planta* and/or disease symptom development, the activated gene is known as a susceptibility (*S*) gene [10]. Alternatively, in some incompatible interactions, TAL effectors may activate an “executor” resistance gene leading to successful host defense [11].

Among the few *S* genes that have been discovered to date, a well-studied case is the *SWEET* gene family of sugar transporters. With one recently described exception [12], induction of any of several clade-3 *SWEET* genes is essential for BLB [13, 14]. It has been suggested that induction of these genes drives accumulation of sugars in the xylem thus providing nutrients for the bacteria, though other possibilities, including osmotic protection for the bacteria or signaling might be relevant outcome. At least three members of the *SWEET* family in rice are direct targets for several *Xoo* TAL effectors, including *OsSWEET11* induced by PthXo1, *OsSWEET13* induced by PthXo2, and *OsSWEET14*, which is induced by multiple TAL effectors from strains of various geographic origins and diverse genetic lineages [10]. Other known virulence targets in rice, although contributing to a lesser extent to host susceptibility, are *OsTFIIA*γ*1* and *OsTFX1*, which respectively encode a general transcription factor subunit induced by PthXo7 and a bZIP transcription factor induced by PthXo6 [15-17]. Interestingly, *OsTFX1* is induced by all *Xoo* strains assessed to date, but its role in promoting disease is not yet understood. *S* genes targeted by TAL effectors from other *Xanthomonas* include *UPA20*, which is responsible for mesophyll cell hypertrophy in bacterial spot of pepper [18], *CsLOB1*, which is involved in pustule formation during citrus bacterial canker [19], *OsSULTR3;6*, which encodes a sulfate transporter important in BLS of rice [20], and a gene encoding a bHLH transcription factor that upregulates a pectate lyase in bacterial spot of tomato [21].

Some TAL effectors act as avirulence factors. Most of them trigger defense responses through the induction of so-called executor (*E*) resistance genes like *Xa27*, *Xa23* and *Xa10* in rice [22-24] or *Bs3* [25] and *Bs4c* in pepper [26]. In *Xoo*, all *avr* genes known so far encode TAL effectors, further underscoring the importance of these proteins in BLB. Some TAL effectors act either as virulence or as avirulence factors, depending on the host genotype. An example is AvrXa7, which induces the *S* gene *OsSWEET14* [27] but also triggers resistance in rice cultivars harboring the executor *R* gene *Xa7* [28]. Another type of resistance triggered by TAL effectors was reported recently involving the recognition of *Xanthomonas oryzae* TAL effectors largely non-specifically by the nucleotide-binding leucine-rich repeat rice Xa1 protein [29] or by a yet unidentified protein encoded at the *Xo1* locus [30].

TAL effectors bind DNA in a specific way owing to their central repeat region, which is composed of almost identical tandem repeats (typically ranging from 33 to 35 amino acids), in which residues at positions 12 and 13 are hypervariable (referred to as repeat variable di-residues, i.e. RVDs) and together dictate a specific interaction with a single nucleotide of the target DNA sequence. Hence, successive RVDs in a TAL effector protein are involved in specific attachment to a sequence of contiguous nucleotides (called the ‘effector binding element,’ EBE) located in the promoter of the gene to be activated [31, 32]. This code-like mechanism underlying TAL effector-DNA specificity enables prediction of TAL effector binding sites in plant genomes [9, 33].

Characterized African strains of *Xoo* have a reduced number of TAL effectors (usually nine) [5], when compared to strains of *Xoc* (up to 28) [34, 35] and Asian *Xoo* (up to 19) [36, 37]. By contrast, weakly virulent US *X. oryzae* strains are devoid of TAL effector genes [38], a characteristic that can be exploited as a genetic tool to study the function of TAL effectors individually [39].

In previous work, we identified two TAL effectors from African *Xoo* strains that play a role in pathogen virulence by targeting the promoter of *SWEET14* at two different EBEs: TalC from the Burkina Faso strain BAI3 [40] and Tal5 from the Malian strain MAI1 [14]. *SWEET14* is the only *S* gene so far known to be targeted by TAL effectors of African strains, as other African *Xoo* TAL effectors have not yet been characterized.

In this study, we employed long-read, single molecule real-time (SMRT) sequencing technology to define the complete TAL effector repertoires (‘TALomes’) of the two African *Xoo* strains MAI1, and BAI3, as has been done recently for a number of *Xo* strains [34, 36, 41, 42]. We also sequenced the genome of strain CFBP1947 [5], from Cameroon, which has been sequenced also by another group [43]. These sequences were used to gain insight into the phylogeny of the strains and the evolution of their TALomes. Further, to evaluate the relative virulence function of each TAL effector within a TALome, we assayed the ability of each to increase the virulence of the TAL effector-less *X. oryzae* strain X11-5A. This strain is less virulent than *Xoo* strains, and the introduction of *Xoo* TAL effectors that induce major *S* genes increases symptom development and pathogen growth *in planta* [39]. X11-5A is therefore a useful tool for dissecting the relative virulence contributions of different TAL effectors. This analysis highlighted TalB as a new, major virulence factor. A search for host genes induced *in planta* by TalB identified the transcription factor genes *OsTFX1* and *OsERF#123* as dual virulence targets of TalB. Activation by designer TAL effectors (dTALEs) delivered by X11-5A or a *talB* knockout derivative of African *Xoo* strain BAI3 revealed *OsERF#123* to be a new class of *S* gene. Our results provide new insights into the diversified strategies underlying the roles of TAL effectors in promoting plant disease.

## Results

### Whole-genome sequence analysis of three African *X. oryzae* pv. *oryzae* strains

SMRT whole genome sequencing was performed for the African *Xoo* strains MAI1, BAI3 and CFBP1947 isolated from Mali, Burkina Faso and Cameroon, respectively [5]. The genomes were assembled using HGAP.3, resulting in one chromosome per strain. The CFBP1947 sequence agreed with the previously published one [accession number CP013666, 43], and whole genome alignments of all three strains revealed only a small (434 kb) inversion in the BAI3 and MAI1 genomes relative to the CFBP1947 genome, yet several rearrangements relative to representative Asian *Xoc* and *Xoo* whole genome sequences, with overall greater structural similarity to the *Xoc* genomes (Fig. 1B, Fig S1A, and Fig S1B). These alignments also revealed the genomes of Asian *Xoo* strains to contain more repetitive sequences overall than *Xoc* or African *Xoo* (Fig S1A).

**Fig 1.**
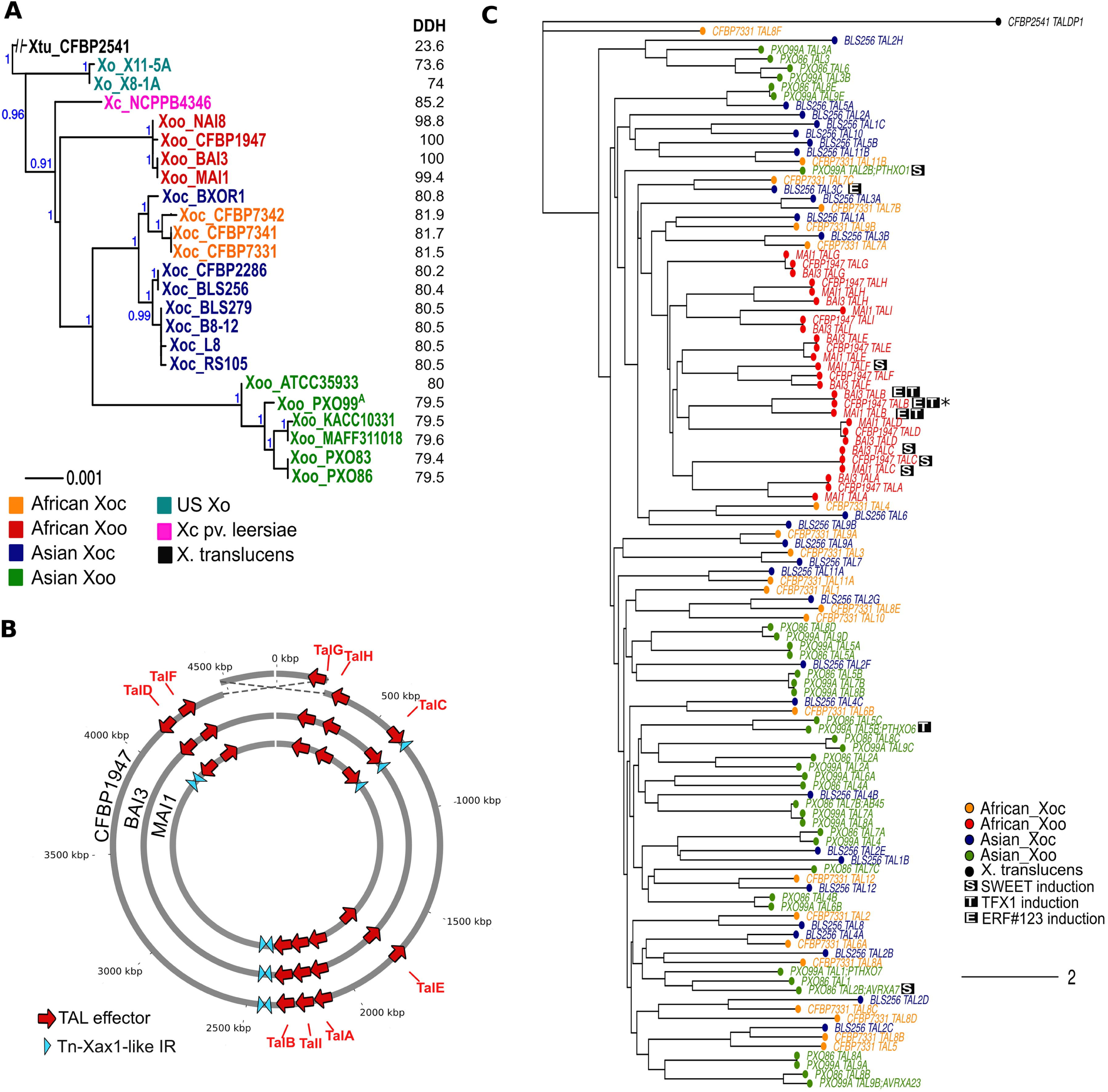
Genomes and TALomes of African *Xoo* compared with other *X. oryzae* strains. **A.** MLSA-based phylogenetic tree of *X. oryzae* strains, built based on alignment of the amino acid sequences of 33 concatenated housekeeping genes using MUSCLE v.3.8.31 [66] and PhyML v. 3.1 [68]. Numbers in blue indicate branch support as calculated by approximate likelihood ratio tests (aLRT) in PhyML v. 3.1 [68]. The *X. campestris* pv. *leersiae*strain NCPPB4346 was included with the *X. oryzae* strains examined. *X. translucens* strain CFBP2541 was used as an out-group (left). Numbers in black indicate in silico DNA-DNA hybridization (DDH) as calculated using Genome-to-Genome Distance Calculator [45] with the *X. oryzae* pv. *oryzae* strain BAI3 as a reference. **B.** Circular representation of three African *X. oryzae* pv. *oryzae* strains showing positions of TAL effector genes and Tn*Xax1*-like inverted repeats. Features are not drawn to scale and position is approximate. Dashed lines highlight a 434kb inversion containing the replication origin in *Xoo* strain CFBP1947 in comparison to strains BAI3 and MAI1. **C.** Neighbour-joining tree built using the program DisTAL [47] based on alignments of repeat regions from TAL effectors from selected *X. oryzae* strains with fully sequenced TALomes. Strains used were (number of TAL effectors in parentheses): *Xoc* = BLS256 (28), CFBP7331 (22), *Xoo*= BAI3 (9), CFBP1947 (9), MAI1 (9), PXO83 (18), PXO86 (18) and PXO99^A^(18). Each tip represents a TAL effector. Tip colors indicate the pathovar and geographic group of the strain; TalDP1 from *X. translucens* CFBP2541 was used as outgroup (accession number WP 039006168.1). S, T and E indicate that the TAL effectors can induce *OsSWEET11 or OsSWEET14*, *OsTFX1* or *OsERF#123* respectively. Asterisks indicate instances where the induction of the corresponding target is hypothesized. Scale bar represents branch lengths as calculated by the nj function of the package ape [69] on a distance matrix of alignments generated by DisTAL [47].

Phylogenetic trees based on multi-locus sequence alignment (MLSA) using 33 housekeeping genes were constructed using available fully sequenced or draft *X. oryzae* genomes, revealing that the three African *Xoo* strains form a closely related group along with the *Xoo* strain NAI8 (draft genome, accession number AYSX00000000.1) collected in Niger [5] (Fig 1A). This analysis also showed that, as previously reported [5, 6], African *Xoo* strains are genetically distant from Asian *Xoo* and appear more closely related to *Xoc*. Also, the *X. campestris* pv. *leersiae* strain NCPPB4346, a pathogen of southern cutgrass shown to cluster with *X. oryzae* strains [38], was revealed to be closely related to both *Xoc* and African *Xoo*. Analyses of Average Nucleotide Identity (Fig S1C) [44] and in-silico DNA-DNA Hybridization [45] support the groupings found using MLSA (Fig 1A). African *Xoo* strains were also found to have, like *Xoc*, an overall reduced number of insertion sequences (IS) elements (Fig S2), and to contain very few (3 to 5) Tn*Xax1*-like short inverted repeats (Fig 1B), when compared to Asian *Xoo*. This particular type of IS which has been found in association with TAL effectors in *Xoo* as well as in other *Xanthomonas*, is hypothesized to be involved in the horizontal transfer of TAL effector genes and their transposition within a genome [46].

**Fig 2.**
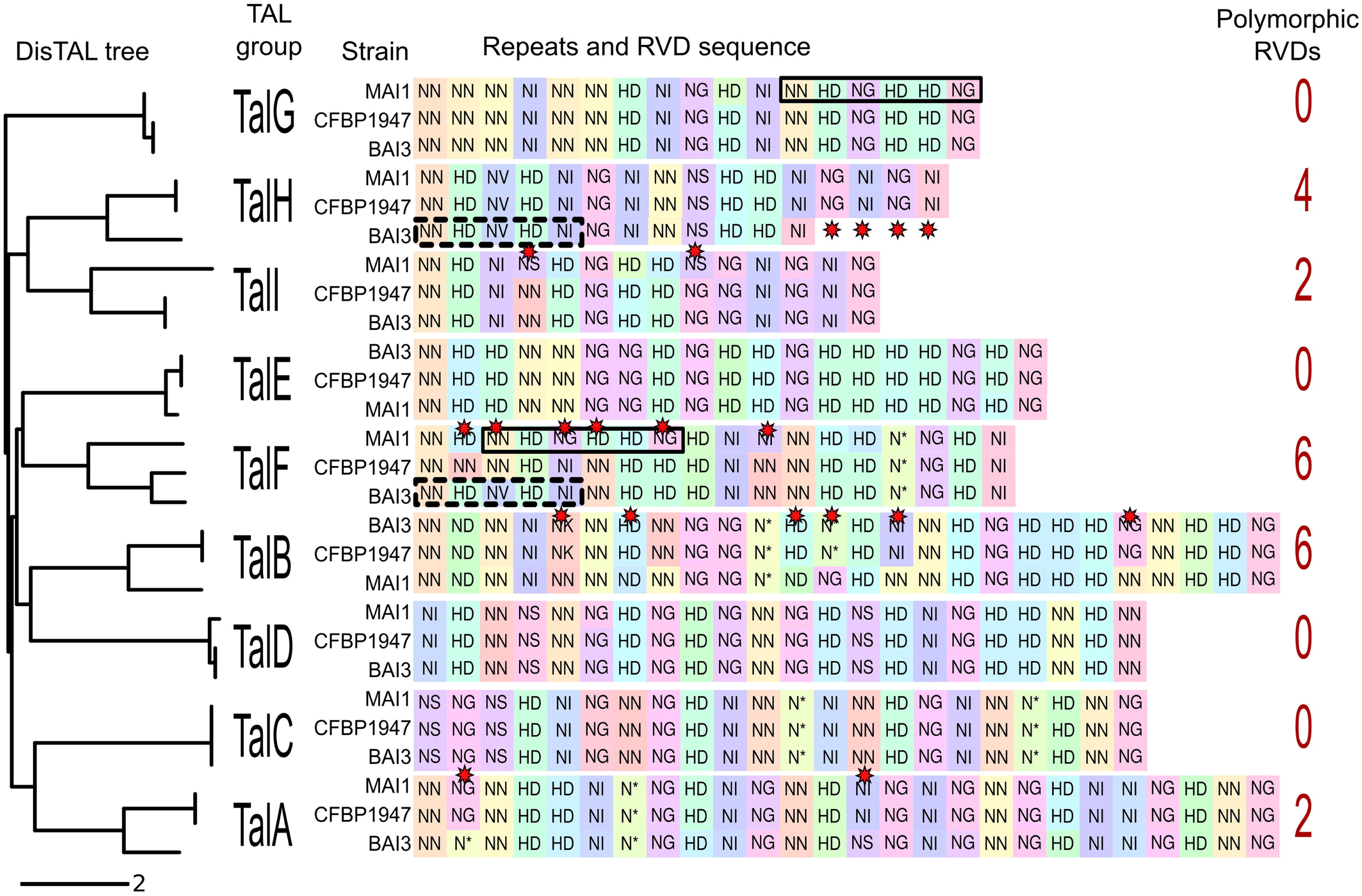
Variation in central repeat sequences and RVDs among African *X. oryzae* pv. *oryzae* TAL effectors. Neighbour-joining tree built using the program DisTAL based on alignment of repeats of TAL effectors from African *Xoo* strains BAI3, MAI1 and CFBP1947. Nine groups are shown and correspond to nine genomic loci (left). RVD sequences are shown for each TAL effector variant from each strain. Background colors for the RVDs represent unique nucleotide repeat sequences, and similar colors indicate similar repeat sequences at the nucleotide level as determined by Needleman-Wunsch alignment. Red stars indicate RVD polymorphisms within one locus. Solid-line rectangles highlight the repeats shared by TalF _MAI1_ and TalG_MAI1_. Dashed-line rectangles highlight those in common to TalF _BAI3_ and TalH_BAI3_.

### Comparative analysis of African *X. oryzae* pv. *oryzae* TALomes and candidate targets

*tal* gene sequences were extracted from the genomes using in-house perl scripts. This identified nine TAL effectors in each strain, including the previously characterized TAL effectors TalC from BAI3 [40] and Tal5 from MAI1 [14]. We used the program DisTAL [47] to assess similarities between the African *Xoo* TAL effectors and TAL effectors from all other fully sequenced *Xoo* and *Xoc* strains by alignment of the RVD sequences. A tree based on these alignments shows that the African *Xoo* TAL effectors group together with TAL effectors from Asian and African *Xoc* strains, suggesting a common ancestral origin (Fig 1C).

Consistent with the fact that the few observed Tn*Xax1*-like inverted repeat sequences discussed above do not flank any TAL effector genes on both sides, across strains the genomic positions of the loci corresponding to the nine TAL effector genes are conserved when accounting for the inversion around the replication origin (Fig 1B). And, the TAL effector gene(s) at each locus across strains are clearly orthologous to one another, falling into nine groups based on DisTAL alignments (Fig 1C). Each locus or TAL effector group is thus identified in this paper with a unique letter, assigned based on the number of repeats, from longest (A) to shortest (I). Specific alleles for each *tal* group are indicated with the strain name in subscript. Tal5 from MAI1 which contains 17.5 repeats was consequently renamed TalF_MAI1_. Alternative names for these effectors following the nomenclatures proposed by others [36, 37] are included in Table S1.

**Table 1.**
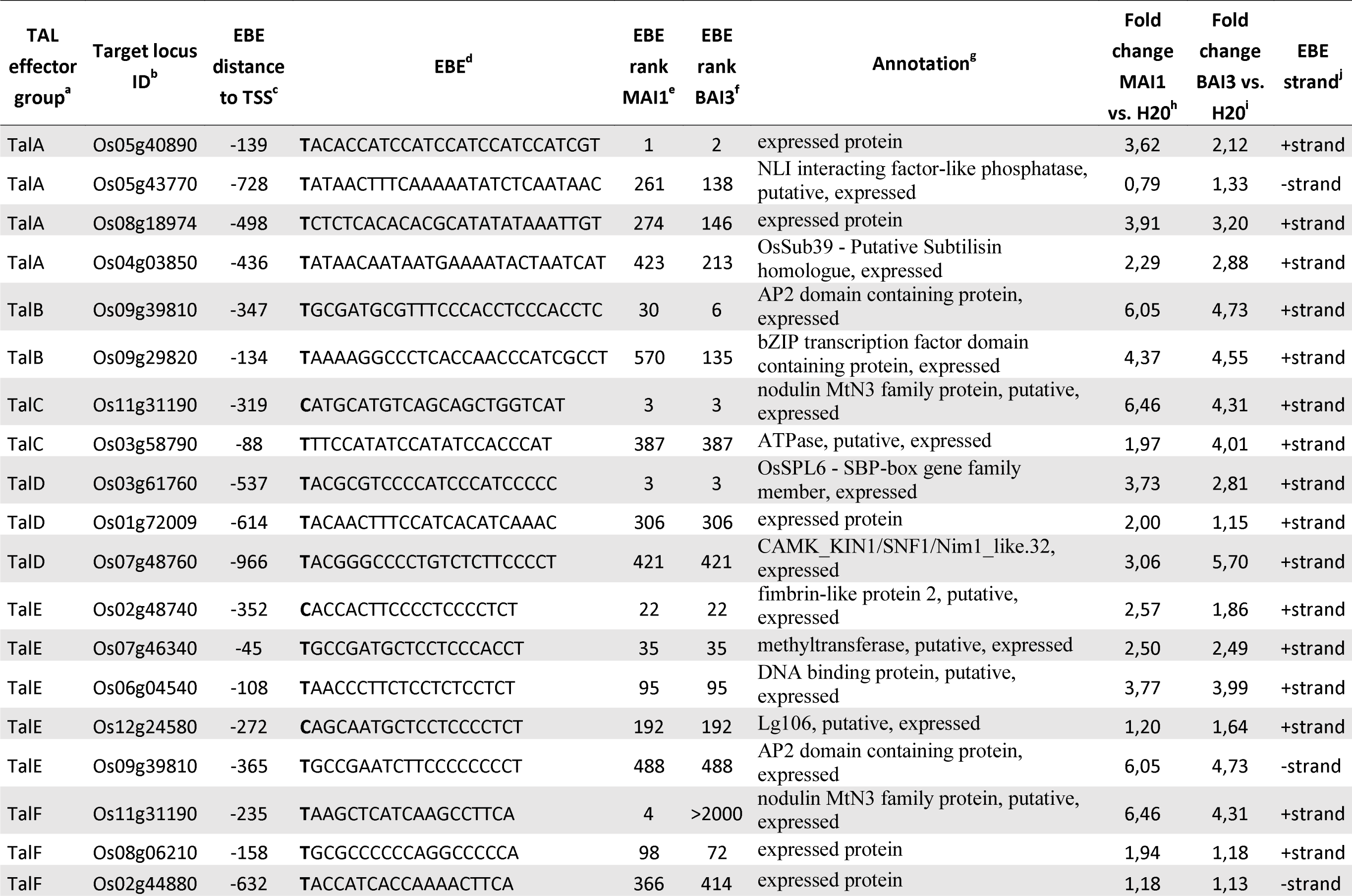
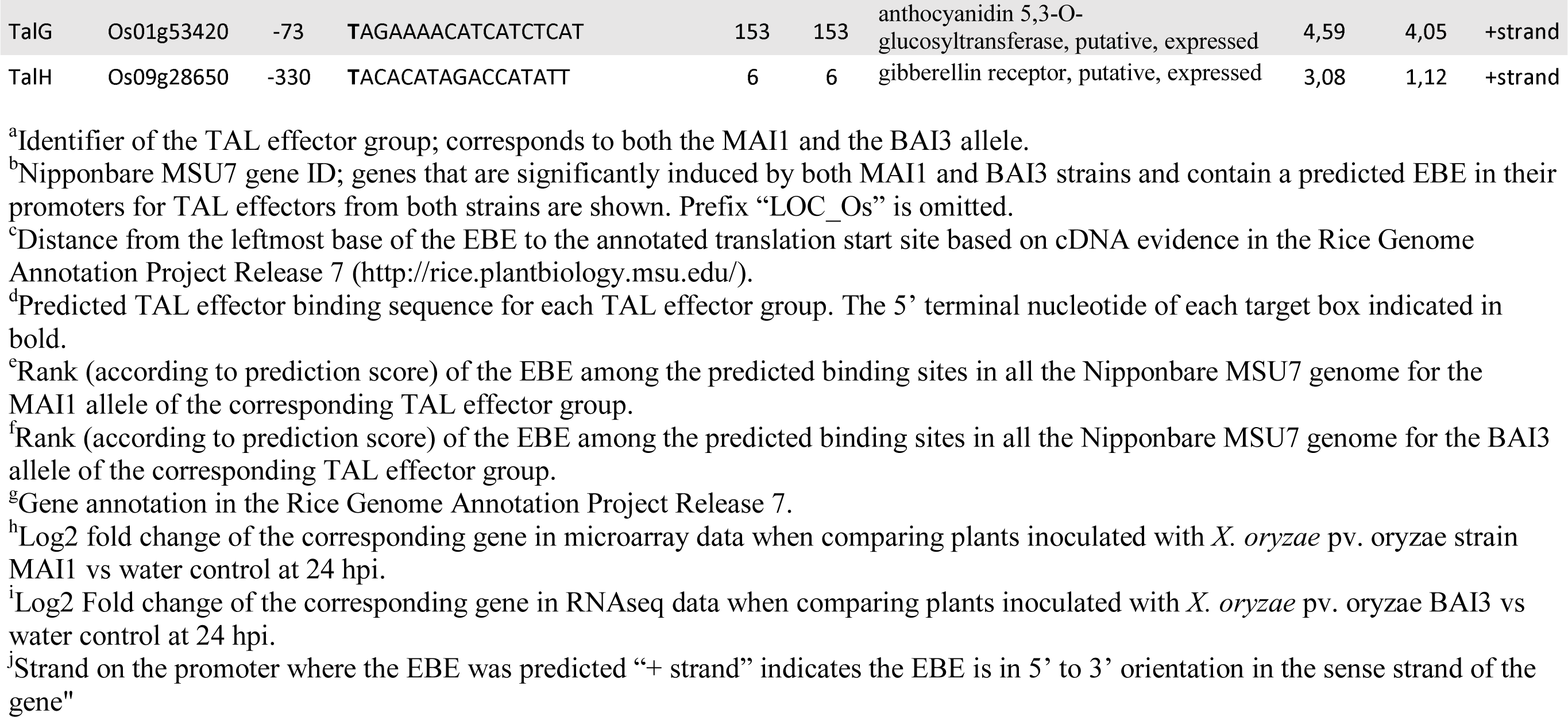
Common targets in the rice genome for TAL effectors from strains MAil and BAI3.

The repeat and RVD sequences are highly conserved across half of the African *Xoo tal* loci, with up to 100% identity at the nucleotide level (e.g., group TalC) (Fig 2). However some loci display variation at the RVD level, particularly the groups TalA, TalB, TalF and TalI, which contain respectively two, six, six, and two polymorphic RVD positions, and the TalH group, where a four-repeat deletion is observed in strain BAI3 (Fig 2). Some of the groups display evidence of recombination. For example, the DNA sequences encoding repeats 3 to 8 of TalF_MAI1_ are different from those in TalF_BAI3_ and TalF_CFBP1947_ but identical to repeats 12 to 17 of TalG_MAI1_ (Fig 2, highlighted in thick frames). Similarly, repeats 1 to 5 of TalF_BAI3_ are identical to those of TalH_BAI3_ (Fig 2, dashed frames), thus suggesting recombination between the TalF and TalG loci in MAI1, and between TalF and TalH in BAI3.

For each of the TAL effectors we predicted EBEs in the rice (cv. Nipponbare) promoterome and looked at the overlap between the predictions for each TAL effector (Fig S3). Based on this analysis, the RVD differences for the polymorphic TAL groups TalA, TalB, TalF, TalH and TaIl can be expected to lead to differences in the activation of some targets. For example, only 60% of the predicted targets are shared between TalA_MAI1_ and TalA_BAI3_ or TalA_CFBP1947_ due to the two RVD polymorphisms found (Fig 2). The differences were more dramatic in the highly variable TalF group where less than 10% of predicted targets for TalF_MAI1_ overlap with those of TalF_BAI3_or TalF_CFBP1947_ (Fig S3).

**Fig 3.**
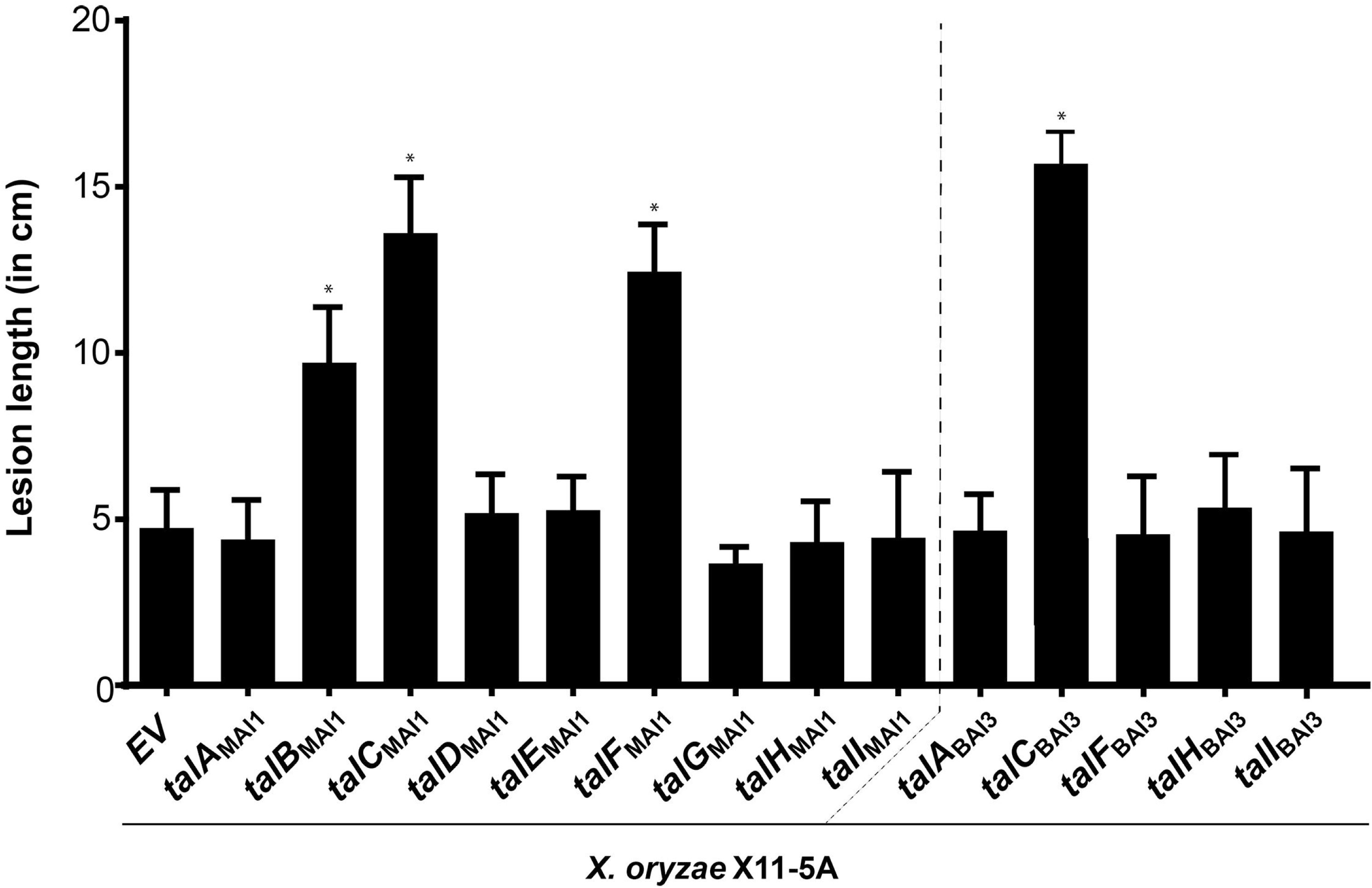
Virulence function of African *X. oryzae* pv. *oryzae* TAL effectors delivered by *X. oryzae* X11-5A. Mean lesion length was measured 15 days post leaf-clip inoculation of the rice variety Azucena with *X. oryzae* X11-5A derivatives carrying the empty vector (EV), p*talC* _BAI3_ used as positive control, or each of the nine individual TAL effector genes from MAI1 and four of those from BAI3 that display RVD polymorphisms relative to their MAI1 counterparts. An asterisk denotes a significant difference from X11-5A(EV) (P < 0.0001). Values represent the averages of at least 10 inoculated leaves. This experiment was repeated four times with similar results.

We next compared the prediction results with expression data. For this, RNA-seq libraries were generated from rice leaves harvested 24 hours after inoculation with the strain MAI1 or a water control. The reads were mapped to the rice genome, and differential expression was determined using the Tuxedo suite [48-50]. 1366 genes were found to be significantly induced in response to strain MAI1. Of these, 152 genes had predicted EBEs in their promoters for MAI1 TAL effectors and were considered candidate TAL effector targets (Table S2). In the case of the strain BAI3, micro-array data were already available for rice plants inoculated with this strain compared to a water control, collected at 24 hours post inoculation (GEO accession GSE19844) [40]. Out of 377 genes induced in response to BAI3, 45 contained predicted EBEs for BAI3 TAL effectors and were therefore considered candidate TAL effector targets (Table S3). We then compared the lists of TAL effector targets for both strains and found that, as shown in Table 1, 20 pairs of TAL effectors and targets were shared between strains BAI3 and MAI1, including *OsSWEET14* being targeted by both TalC_MAI1_ and TalC_BAI3_.

### Systematic functional analysis of African *Xoo* TALomes

We reasoned that the list of shared targets described above may include novel *S* genes that contribute to the severity of disease caused by each strain. As a first step to identify such genes, we subcloned the TALomes of strains BAI3 and MAI1 to test for virulence contributions by gain of function assays using X11-5A, the *Xanthomonas oryzae* strain that is naturally devoid of TAL effector genes [38]. A strategy based on the enrichment of *Bam*HI-TAL effector sequences from genomic DNA libraries [51], led to the isolation of all nine *tal* genes from strain MAI1 and eight from strain BAI3 (we were not able to isolate talB_BAI3_ despite several attempts). The sequences of the cloned genes matched those from the genome assembly.

For functional assays, we aimed at testing TAL effectors representing most of the TALome diversity found in African *Xoo* so far. To that end we selected the nine TAL effectors from MAI1, four from BAI3 that displayed variations at the RVD level with respect to their counterpart in MAI1 (namely TalA_BAI3_, TalF_BAI3_, TalH_BAI3_ and TalI_BAI3_, Fig 2), and, for reference and as a positive control the previously characterized major virulence factor TalC_BAI3_ [40]. Each *tal* gene was conjugated individually into X11-5A and the resulting strains were assayed by leaf clip inoculation to leaves of the susceptible rice variety Azucena for increases in virulence (lesion length after 15 days) relative to an empty vector conjugant. Three of the nine MAI1 TAL effectors, namely TalB_MAI1_, TalC_MAI1_ and TalF_MAI1_, enhanced X11-5A virulence (Fig 3). TalC_MAI1_ and TalF_MAI1_ are discussed below, and TalB_MAI1_ is discussed in the following section.

The RVD array of TalC_MAI1_ is 100% identical to that of TalC_BAI3_ (Fig 2), which activates the expression of the *S* gene *OsSWEET14* [40], thus explaining its ability to increase the virulence of X11-5A. TalF_MAI1_ was shown previously also to induce *OsSWEET14* by binding at a non-overlapping EBE [14], likewise explaining its ability to enhance X11-5A virulence. To our knowledge, this is the first example of a strain carrying two TAL effectors that target the same *S* gene promoter at unrelated EBEs, and may reflect a pathogen adaptation to overcome host loss-of-susceptibility alleles such as *xa41*(t) that disrupt an EBE [52]. In contrast to TalF_MAI1_, TalF_BAI3_ did not increase the virulence of X11-5A (Fig 3). Indeed no EBE is predicted in the *SWEET14* promoter for TalF_BAI3_, which as noted above differs at 6 out of 18 RVDs from TalF_MAI1_.

### TalB_MAI1_ contributes to virulence by targeting both a known *S* gene and a new one

To understand how TalB_MAI1_ promotes disease, we sought to identify its relevant targets. Among the nine genes induced by BAI3 that each has a predicted TalB EBE in its promoter (Table S3), we focused on two that were also induced by MAI1 (Table S2). *Os09g39810,* which was the most highly induced by MAI1 and had the highest TalB_MAI1_ EBE prediction score (Table 1, Table S2), and *Os09g29820* (*OsTFX1*), which was previously reported as a BLB *S* gene [15, 16]. *Os09g39810* is annotated as an expressed protein containing an AP2 domain and is from here on referred to as *OsERF#123* [annotated as an expressed protein containing an AP2 domain, 53]. Both TalB_MAI1_ and TalB_BAI3_, despite their six polymorphic RVDs, are predicted with high scores to bind to the same EBE in its promoter. This EBE ranks number 30 in the genome for TalB_MAI1_ and number 6 for TalB_BAI3_ (Table 1, Table S2 and Table S3). *OsTFX1* encodes a bZIP transcription factor, it is highly induced upon infection of rice with MAI1, and its predicted EBE ranks 570 in the genome for TalB_MAI1_ and 135 for TalB_BAI3_.

To ascertain whether *OsERF#123* and *OsTFX1* are bona fide targets of TalB_MAI1_, rice leaves were infiltrated with MAI1 and X11-5A(p*talB*_MAI1_), respectively carrying *talB*_MAI1_ on the chromosome or on a plasmid, and X11-5A carrying an empty vector. As expected, RT-PCR amplification showed that both *OsERF#123* and *OsTFX1* were highly and specifically induced upon infection of rice with the strains expressing TalB_MAI1_ (Fig S4). With the aim of deciphering which of the two targets is biologically relevant, i.e. an *S* gene, we engineered dTALEs to specifically induce each of them individually. Two dTALEs per candidate gene were devised, targeting sites located in the vicinity of the native EBE (Fig 4A). Leaf-clipping inoculation assays using X11-5A derivatives showed that dTALE-mediated induction of *OsERF#123* enhanced virulence similarly to TalB_MAI1_ (Fig 4B). dTALEs for *OsTFX1* also conferred higher virulence to X11-5A albeit to a lower extent than the *OsERF#123* dTALEs. The virulence contribution was similar to that conferred on X11-5A by TAL effector PthXo6 from the Asian *Xoo* strain PXO99^A^ [15, 16], which was included as a positive control. Importantly, X11-5A strains expressing the *OsERF#123*-targeted dTALEs, dTALE_ERF-1_ or dTALE_ERF-2,_ induced *OsERF#123* and not *OsTFX1*, as shown by qRT-PCR analysis (Fig 4C). Last, we evaluated patterns of *OsERF#123* transcript initiation using 5’-RACE experiments, and discovered that TalB initiates transcription primarily at sites residing 41-69 bp downstream of the 3’ end of its EBE (Fig S5), as is the case for many TAL effectors and their targets. Altogether, our results reveal that TalB_MAI1_ is a virulence factor that activates two *S* genes, *OsTFX1* and *OsERF#123*, each of which functions independently.

**Fig 4.**
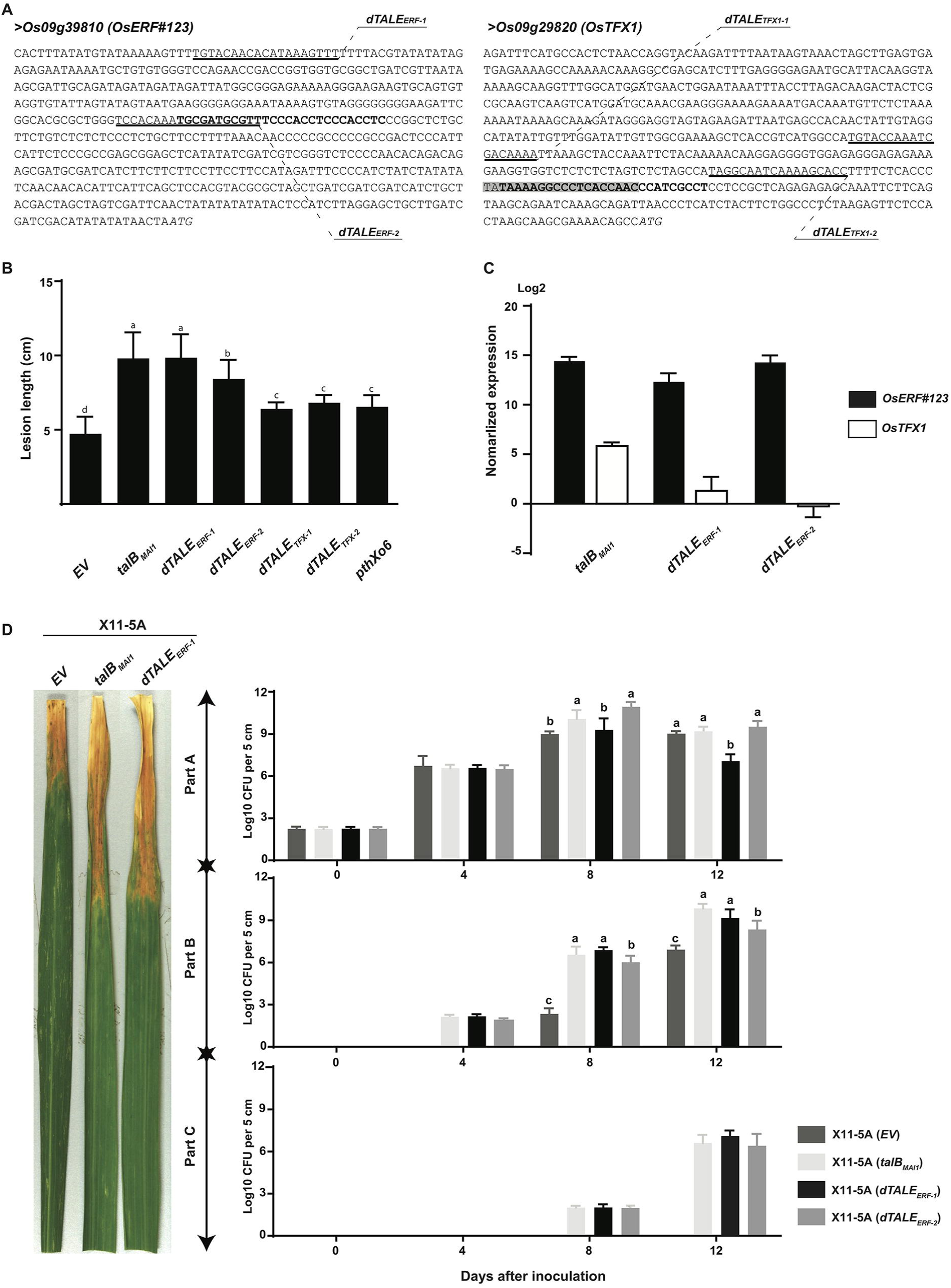
TalB_MAI1_ targets *OsTFX1* and a novel *S* gene *OsERF#123*. (A) DNA sequence of the promoter regions of TalB_MAI1_-induced genes *Os09g39810*(aka *OsERF#123*) and *Os09g29820*(aka *OsTFX1*) in rice cv. Nipponbare. The effector binding elements (EBEs) for TalB_MAI1_ and PthXo6 from the Asian *Xoo* strain PXO99^A^are depicted in bold and highlighted in gray, respectively. The EBEs for gene-specific dTALEs used in panels B and C are underlined and labeled. Italics indicate translational start sites, based on the rice Nipponbare v. MSU7 genome (http://rice.plantbiology.msu.edu/). (B) dTALEs targeted to *OsERF#123* and dTALEs targeted to *OsTFX1* increase the virulence of Xo strain X11-5A. Rice leaves of Azucena were inoculated by leaf-clipping with derivatives of X11-5A carrying an empty vector (*EV*) or expressing talB_MAI1_, dTALE_ERF-1_, dTALE_ERF-2_, dTALE_TFX-1_ dTALE_TFX-2_ *o* or *pthXo6*. Lesion length was measured at 15 days after inoculation (dai). Data are the mean of at least eight measurements. Error bars represent +/-SD. Bars with same letters are not statistically different based on a Tukey’s multiple comparisons test (α = 0.05). Data were plotted and statistical analyses were performed using the Graphpad prism 6 software. (C) dTALE_ERF-1_ and dTALE _ERF-2_ specifically activate *OsERF#123* but not *OsTFX1*. The expression levels of *OsERF#123* and *OsTFX1* were evaluated by quantitative reverse transcription polymerase chain reaction (qRT-PCR) 24 hours after syringe infiltration of Azucena rice leaves with X11-5A strains carrying an empty vector (*EV*) or derivatives expressing talB_MAI1_, dTALE_ERF-1_ *o* or dTALE_ERF-2_. Gene expression was normalized against X11-5A (*EV*). Error bars represent +/-SD. (D) Population size and distribution of X11-5A(*EV*), X11-5A(ptalB_MAI1_), X11-5A(pdTALE_ERF-1_) and X11-5A(pdTALE_ERF-2_) in leaves of cultivar Azucena following leaf-clipping inoculation. Four-week-old plants were inoculated and bacterial populations in 5cm leaf segments were measured at 0, 4, 8, and 12 days after inoculation. Values represent averages of three inoculations. The experiment was repeated three times with similar results. Error bars represent +/-SD. Bars with the same letters are not statistically different, based on a Tukey’s multiple comparisons test (α = 0.05). Data were plotted and statistical analyses were performed using the Graphpad prism 6 software. Left panel: representative lesions photographed at 14 dai. This experiment was reproduced three times with similar results.

**Fig 5.**
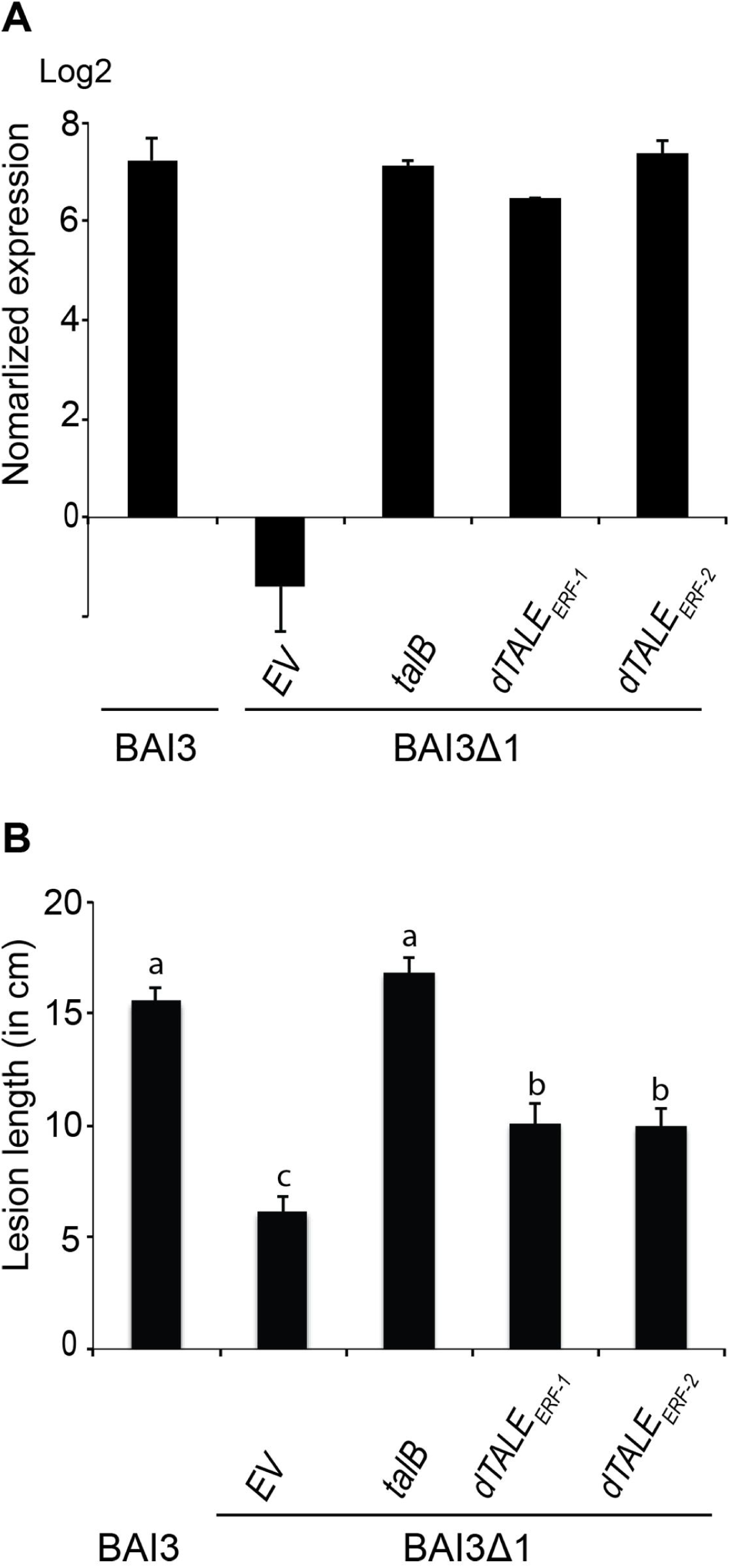
Mutation of *talB* in BAI3 results in a decrease of virulence partially complemented by the induction of *OsERF#123*. (A) BAI3 mutant strain BAI3Δ1 carrying plasmid-borne talB_MAI1_, dTALE_ERF-1_ *o* or dTALE_ERF-2_ *b* but not BAI3Δ1 carrying the empty vector (*EV*) induces *OsERF#123*. The expression level of *OsERF#123* was evaluated by quantitative reverse transcription polymerase chain reaction (qRT-PCR) 24 hours after syringe infiltration of Nipponbare rice leaves. The wild type strain BAI3 was used as control. Gene expression was normalized against water inoculated leaves. Error bars represent +/-SD. (B) TalB and dTALEs _ERF_ act as virulence factors for *Xoo* strain BAI3. Rice leaves of IR24 were inoculated by leaf-clipping with BAI3Δ1 strains carrying an empty vector (*EV*) or derivatives carrying talB_MAI1_, dTALE_ERF-1_, or dTALE_ERF-2_. Lesion length was measured at 15 days after inoculation (dai). Data are the mean of at least 16 measurements. Error bars represent +/-SE. Bars with same letters are not statistically different based on a Tukey’s multiple comparisons test (α = 0.05).

To confirm the function of TalB as a virulence factor for African *Xoo*, we generated a library of BAI3 TAL effector gene mutants by transformation with the suicide plasmid pSM7 [20] and identified a *talB* mutant by screening for strains unable to induce *OsERF#123*, assessed by qRT-PCR 24h after infiltration (data not shown). We identified such a strain and designated it BAI3Δ1 (Fig 5A). Assayed by leaf clip inoculation, BAI3Δ1 showed reduced virulence relative to the wildtype strain (Fig 5B). Induction of *OsERF#123* and virulence were fully restored by plasmid-borne *talB*_MAI1_, while induction of OsERF#123 was fully restored and virulence partially restored by either of the two *OsERF#123*-targeted dTALEs (Fig 5A and Fig 5B), thus establishing that TalB contributes to virulence in its native context by activating the newly identified *S* gene *OsERF#123*.

## Discussion

In this study we sequenced the genomes of three African strains of *Xoo* and characterized their TALomes systematically using a gain-of-virulence assay, providing a first glimpse into the function and evolution of TAL effectors in this particular group of *X. oryzae* strains. Based on RNA profiling and *in silico* prediction, and both gain and loss of virulence assays and targeted dTALEs, we identified TalB as a virulence factor and TalB target *OsERF#123* as a novel *S* gene for bacterial blight of rice.

Our analysis revealed each African *Xoo* strain to contain nine TAL effectors that can be classified according to their repeat and RVD sequences [36, 47] in nine groups, corresponding to nine syntenic loci. Five of these loci showed some degree of variation in RVD sequence between strains, variation that can translate into different effects on the plant transcriptome. When comparing these sequences with the results of inoculation assays (see below), no clear relation was seen between contribution to virulence and evolutionary conservation. Among the three TAL effectors from MAI1 that contributed to virulence, TalC_MAI1_ is highly conserved (i.e. identical at the nucleotide level) in all three strains and is expected to contribute to virulence in the same way in all strains harboring it, by inducing *OsSWEET14*, as previously reported [40]. Meanwhile, TalB_MAI1_ shows some RVD differences from TalB_BAI3_ and TalB_CFBP1947_; nonetheless, the two latter alleles are expected also to induce the relevant targets *OsTFX1* and *OsERF#123* and thus to contribute to virulence similarly. In the case of TalB_BAI3_ there is transcriptomic (Table 1) and functional (Fig 5) evidence that this is the case. Finally, TalF_MAI1_, which also induces *OsSWEET14*, at an EBE distinct from that of TalC, belongs to a highly variable group in which the alleles in strains BAI3 and CFBP1947 may be inducing completely different targets. And at least in the case of BAI3, no significant contribution to virulence by its allele *talF*_BAI3_ was observed in inoculated Azucena plants (Fig 3). The degree of variation in each of these groups may be dictated by interaction with the host, specifically the degree of variation at the target promoters across different host genotypes.

The evolution of the *talF* locus is of particular interest. The sequence analysis presented in this work hints at possible recombination events leading to variants of this TAL effector in the strains BAI3 and MAI1, where in each case a set of contiguous repeats was found to match exactly those of another TAL effector in the genome (different in each case). The overall high similarity of TAL effector repeats likely favors recombination, and recombination of repeats may indeed be responsible for fast evolution of TAL effector variants in *Xanthomonas* [46, 47, 54]. Recombination in this case may have been further favored by the genomic context given that the loci involved are found close to the replication origin (Fig 1B), a region known to be a hotspot for recombination [55].

The fact that *talF*_MAI1_ targets the *S* gene *OsSWEET14* convergently with TalC in MAI1 raises questions about the selective forces acting upon this locus. It is possible that the MAI1 variant *talF*_MAI1_ was positively selected after a recombination event allowing for its induction of *OsSWEET14;* indeed, the recombined repeats match the corresponding nucleotides in the EBE almost perfectly according to the TAL code (Table 1, Fig 6). However, how could this selection have occurred in the presence of the *OsSWEET14* inducer TalC_MAI1_ in the same strain? One possibility is that these two TAL effectors have been maintained in the *Xoo* strain MAI1 in response to genomic variants in the host population. Indeed, at least one type of variation (named *xa41*(*t*)) has been described in African rice varieties that impairs binding of TalF_MAI1_ to its EBE in the promoter of *OsSWEET14* [52]. It is possible that a similar loss-of-susceptibility allele that impairs binding of TalC_MAI1_ also exists in rice and is driving the selection of TalF_MAI1_. Alternatively, TalF_MAI1_ may be an ancient TAL effector that was lost in strains BAI3 and CFBP1947 after escaping selection due to its redundancy with TalC. Studying the TALomes of more African *Xoo* strains and genetic variation at *OsSWEET14* will likely clarify the evolutionary history of these loci.

**Fig 6.**
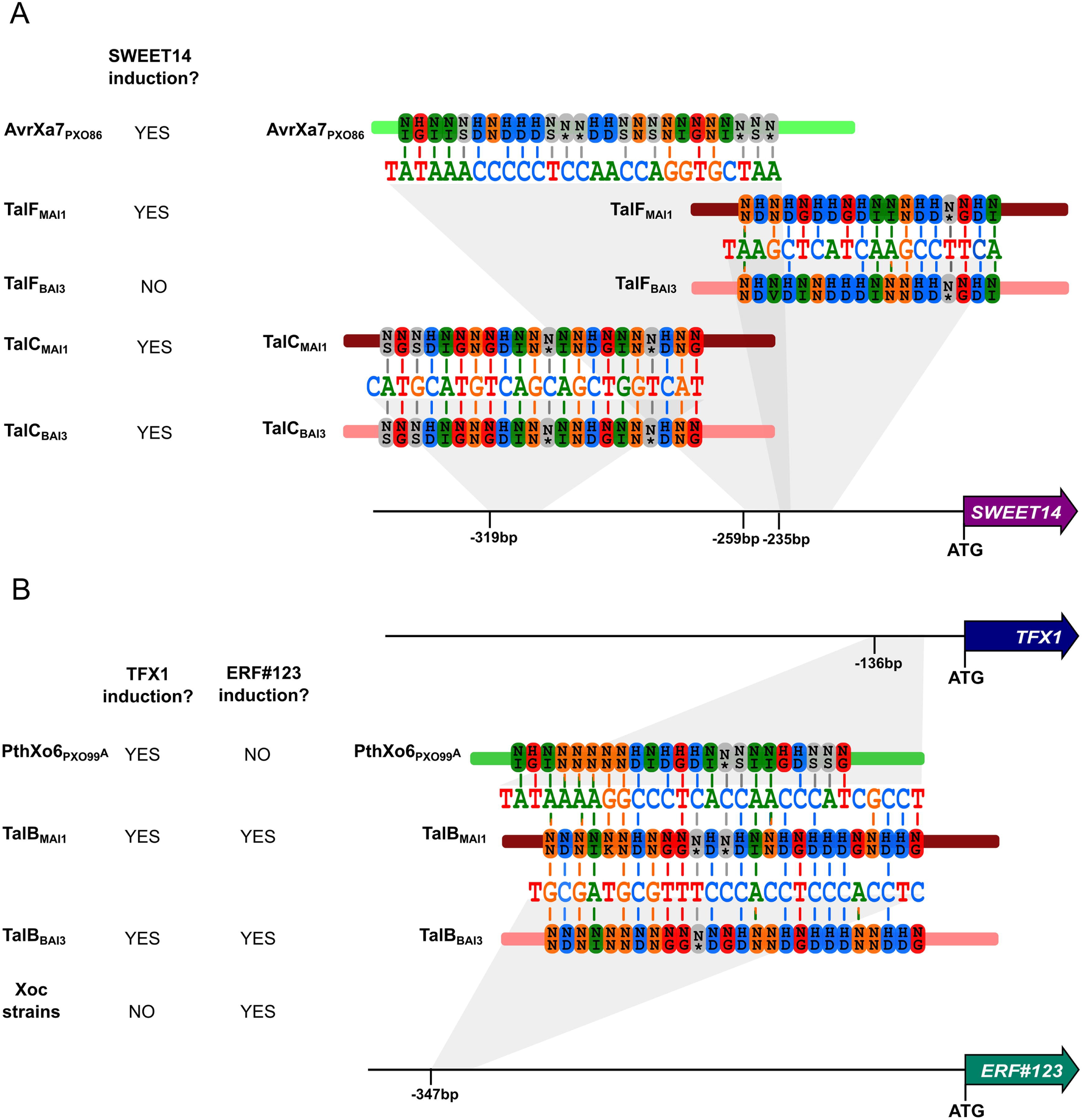
Functional convergence and dual targeting activity among African *X. oryzae* pv. *oryzae* TAL effectors. Targeting of three susceptibility genes by multiple TAL effectors is shown. Not included in the figure, *OsERF#123* is also induced by Tal3c from *Xoc* strain BLS256 upon binding to two EBEs, different from the one bound by TalB [61]. Genes are indicated as arrows, black horizontal lines correspond to the promoter region of the gene, and the distance of the targeted EBEs to the annotated translation start site is shown. EBE sequences are shown color-coded by nucleotide, and RVD sequences of TAL effectors are shown color-coded according to their nucleotide specificity. A vertical line between a nucleotide and an RVD highlights a match. To the left is indicated by YES instances where there is evidence that the corresponding gene is induced by the TAL effector, and by NO instances where there is no such evidence.

With the aim of evaluating the function of the TAL effector repertoires of two African *Xoo* strains, we used the TAL effector-less strain of *X. oryzae* X11-5A [38]. Previous studies had shown that three TAL effectors that contribute to virulence of their strains of origin by targeting clade-III members of the SWEET family of sugar transporters conferred higher virulence to X11-5A [39]. Here, we took advantage of this gain-of-function system to systematically decipher the individual function of each TAL effector within the TALomes of the three African *Xoo* strains. This approach is easier than loss-of-function mutagenesis approaches, and unlike such approaches, it avoids the pitfall of redundancy among effectors. Indeed, the role of *talF*_MAI1_ could not have been identified by loss-of-function screening since *talC*_MAI1_ activates the same *S* gene it does.

Our gain-of-function approach led also to the discovery that TalB_MAI1_ is a novel major virulence factor. Combining *in silico* prediction of EBEs and transcriptomic data, we pinpointed two candidate targets, *OsERF#123* and *OsTFX1*. By specifically inducing the expression of each using dTALEs, we found that each target could explain some of the disease promoting effect of TalB_MAI1_. Interestingly *OsTFX1* had been reported as a *S* gene for Asian *Xoo* strains and is known to be targeted by the TAL effector PthXo6 of *Xoo* strain PXO99^A^ [15, 16]. Our data show that African *Xoo* strains also have acquired the capacity to activate *OsTFX1* albeit with a TAL effector harboring a totally different RVD array (Fig 6). In addition to the well-known example of *OsSWEET14* whose promoter is targeted by several unrelated TAL effectors [56], *OsTFX1* represents a new case of functional convergence wherein two completely unrelated TAL effectors in strains of different geographic origin and distant genetic lineages have evolved to bind to overlapping EBEs. In line with this conclusion, *OsTFX1* is induced by several strains of Asian *Xoo* [15], suggesting that its induction is as essential as the induction of clade-III sugar transporters of the SWEET family. Yet, the regulatory targets of *OsTFX1* and the physiological consequences of its induction remain to be characterized.

The other gene targeted by TalB_MAI1_, *OsERF#123*, encodes a putative protein of 131 amino acids (BAT09530) characterized by an AP2/ERF domain (spanning from residues 16 to 75). BLAST analysis showed that *OsERF#123* is single-copy in the rice cv. Nipponbare genome and broadly conserved across the *Oryza* genus (not shown). It attracted our attention because it was the most highly induced of the candidate TalB_MAI1_ targets, and it had never been described before. In a genome-wide analysis of the ERF family in rice and Arabidopsis, Nakano *et al*. [53] showed that rice ERF proteins sort into 15 groups which can be further divided into subgroups, based on sequence polymorphisms. Interestingly, group IX consists of proteins specialized in mediating disease resistance. OsERF#123 was assigned to the subgroup IXc which includes a few proteins whose biological function is related to the induction of defense responses such as SlPti5, described as a pathogenesis-related (PR) protein [57, 58]. Nevertheless, it cannot be excluded that OsERF#123 acts as a negative regulator of defense responses as is the case for some WRKY transcription factors that tightly regulate other, positive regulatory WRKY factors [59]. In such a scenario, TAL effector-mediated induction of *OsERF#123* could result in increased susceptibility. Strikingly, surveying publicly available expression data, we found evidence of *OsERF#123* expression only in response to African *Xoo* [40], *Xoc* [34], or chilling stress [60]. Semi-quantitative RT-PCR confirmed that a panel of 15 African *Xoo* strains originating from Burkina Faso and Mali induced *OsERF#123*, while the Asian *Xoo* strain PXO99^A^ did not (not shown). Also, examination of RNAseq data generated from rice leaf tissue 48h after inoculation with ten geographically diverse *Xoc* strains [34], revealed that *OsERF#123* was highly induced by nine out of the ten strains. This result suggests that *OsERF#123* might be targeted by strains of *Xoc* to promote bacterial leaf streak and is in agreement with our current understanding that African *Xoo* are closer to *Xoc* than to Asian *Xoo* strains. In separate work, Wang *et al*. [61] have recently shown that Tal3c of the *Xoc* strain BLS256, which is entirely distinct from TalB_MAI1_ in RVD sequence, is responsible for *OsERF#123* induction, highlighting yet another striking case of evolutionary convergence.

Altogether, our results show that despite having a reduced TALome relative to their Asian counterparts or *Xoc,* African *Xoo* strains have successfully adapted that TALome to target physiologically relevant genes. As evidence, we show two features of African *Xoo* TALomes that have not been reported previously. First, we show redundant and convergent targeting of the same *S* gene by two TAL effectors from one strain (MAI1) with different RVD sequences: TalC_MAI1_ and TalF_MAI1_ (Fig 6), a feature that may have arisen as a response to loss-of-susceptibility alleles in the host population [52]. In the case of TalF_MAI1_ we also hint for the first time at a recombination event leading to the evolution of virulence activity.

Second, we describe for the first time a TAL effector, TalB_MAI1_, with dual targeting activity, binding to two different EBEs in unrelated genes, where both targets contribute to susceptibility. While TAL effectors often have been shown to induce multiple genes, so far at most only one of the induced genes has been shown to be required for virulence [19, 20]. Interestingly, in at least one case, the TAL effector may have higher affinity for the physiologically relevant target, based on EBE prediction scores [20]. It is then curious that the BAI3 and CFBP1947 variants of TalB are predicted to bind with even higher affinity to the EBEs of both *OsTFX1* and *OsERF#123* than TalB_MAI1_ is (Table 1), suggesting that the *talB* locus is undergoing some evolutionary fine-tuning of its binding specificity to target both genes simultaneously. Dual targeting of two susceptibility genes has recently been hypothesized for another African *Xoo* TAL effector, TalC, although its secondary target is yet to be discovered [12].

Finally, both TalB targets, *OsTFX1* and *OsERF#123*, similarly to *SWEET* genes may constitute widely exploited hubs for pathogenicity, targeted convergently by distinct TAL effectors of different strains. Unlike *SWEET* genes, as transcription factor genes they may also serve uniquely as regulatory hubs for broad manipulation of the host transcriptome by TAL effectors, although their targets remain to be elucidated. Since *OsERF#123* is induced by African *Xoo* strains and diverse *Xoc* strains, it may be of particular interest as an *S* gene for both bacterial blight and bacterial leaf streak. However its role in bacterial leaf streak has yet to be determined [61]. Along with *OsHen1* (LOC_Os07g06970), *OsERF#123* is so far one of only a few genes targeted both by *Xoc* and by *Xoo* TAL effectors [32, 62, 63]. If it contributes to leaf streak susceptibility it would be the first example of a susceptibility gene for both diseases.

Deciphering the virulence function of individual TAL effectors and identifying the host target(s) is key to discovering or engineering novel sources of resistance to combat bacterial blight. This is particularly true in the case of African *Xoo* strains, which differ in many aspects from their Asian counterparts, and for which sources of resistance effective against Asian strains may not be effective. In this study, we have laid a foundation for such inquiry and discovered both convergent targeting (of *S* genes *OsTFX1* and *OsSWEET14*) by Asian and African *Xoo* strains, as well as a novel susceptibility gene (*OsERF#123*) for bacterial blight that is targeted by African *Xoo* strains and diverse *Xoc* strains, but so far not by Asian *Xoo* strains. Further experiments are in progress to investigate how this novel class of virulence target confers susceptibility to bacterial blight, and whether it plays a role in leaf streak.

## Methods

### Bacterial strains, growth conditions and plasmids

The bacterial strains used in this study were *Escherichia coli* DH5α (Stratagene, La Jolla, CA, USA), *X. oryzae* pv. *oryzae* strains BAI3, MAI1 and CFBP1947 [5], and *X. oryzae* strain X11-5A [38]. *E. coli* cells were cultivated at 37° C in Luria-Bertani broth (LB) medium, and *X. oryzae* strains at 28°C in PSA medium (10 g l^−1^ peptone, 10 g l^−1^ sucrose, 1 g l^−1^ glutamic acid, 16 g l^−1^ agar). Plasmids were introduced into *E. coli* by electroporation and into *Xo* by biparental conjugation using *E. coli* strain S17-1 as previously described [40]. The plasmid used to subclone individual *tal* genes is a derivative of pSKX1-*talC* obtained upon digestion with *Bam*HI and direct self-religation, leading to a construct harboring the ATG start codon of *talC* and 151 3’ bp [14]; expression is driven by the *lac* promoter, which is constitutive in *Xanthomonas*. BAI3Δ1 was generated by transformation of BAI3 with the suicide plasmid pSM7 and mutant strains were selected with kanamycin at 25μg/mL [20].

### Plant material and plant inoculations

Experiments were performed under glasshouse conditions under cycles of 12 hours of light at 28-30°C and 80% relative humidity (RH), and 12 hours of dark at 25°C and 70% RH. The rice varieties Azucena and IR24 were inoculated by leaf-clipping of four-week-old plants and evaluated as reported previously [40]. In each experiment, at least eight leaves from eight individuals were inoculated and experiments were replicated three times independently. Growth curves experiments were achieved as reported previously [40].

### DNA preparation and purification

Total genomic DNA of the three African *Xoo* strains was prepared using the QIAGEN midi-prep genomic DNA preparation protocol for bacteria and using the Genomic-tip 100/G (Qiagen, Valencia, CA, USA). DNA was re-suspended in 500 μl of ddH_2_O and aliquoted into several tubes and stored at -20oC for SMRT sequencing and sub-cloning of *tal* genes. All the plasmid preps were carried out by using the QIAprep Spin Miniprep Kit (Qiagen). DNA purification was performed by using the QIAquick PCR Purification Kit (Qiagen).

### RNA isolation and qRT-PCR

Leaves of 3-week-old rice plants were infiltrated with water or *Xo* strains using an OD_600_ of 0.5. At 1 dpi, leaf segments from three plants were ground into a fine powder using the Qiagen Tissue-Lyser system (30 rps for 30 s). Total RNA was isolated from rice leaves using the RNeasy kit (Qiagen). cDNA was generated as reported previously [14]. Real-time PCR was performed on the Mx3005p multiplex quantitative PCR system (Stratagene, La Jolla, CA, USA) using SYBR Green (Eurogentec, Liège, Belgium). For each gene, three independent biological replicates were analyzed. DNA amounts were normalized using actin as a reference gene. Primer sequences are provided in Table S4.

### RNAseq analysis

RNAseq libraries (paired-end strand specific TRUEseq Illumina) were constructed from RNA extracted from leaf tissues of Nipponbare plants collected 24 hours post inoculation with the *Xoo* strain MAI1 and a water control. Libraries were sequenced in a HiSeq2000 machine (Illumina) at the INRA Unité de Recherche en Génomique Végétale (Evry, France). Library quality was verified using FastQC v0.10.1 (http://www.bioinformatics.babraham.ac.uk/projects/fastqc/). Reads were mapped against the rice genome (Nipponbare v. MSU7) using the Tuxedo suite v2.1.1 [48] with the following parameters: Tophat (--max-multihits 20--b2-very-fast); Cufflinks (--frag-bias-correct --library-type fr-secondstrand --multi-read-correct); Cuffdiff (--upper-quartile-norm --multi-read-correc --FDR 0.05). Differentially expressed genes were determined using the R packages cummerbund [48-50]. Genes identified as differentially expressed (pvalue >0.01) by at least one of these approaches were kept for analyses. Libraries are available under accession GSE108504 at the NCBI-GEO.

### Construction of designer TAL effectors

The *OsERF#123* and *OsTFX1* promoter sequences from the rice variety Nipponbare were analyzed in order to design the appropriate designer TAL effectors (dTALE) binding sites as described previously [14]. The specificity of the selected EBEs were controlled using Talvez v.3.1 (http://bioinfo.mpl.ird.fr/cgi-bin/talvez/talvez.cgi; [62]). dTALEs were generated using the Golden TAL technology and expressed as FLAG fusions under the control of a *lac* promoter in the Golden Gate-compatible broad host range vector pSKX1 [14]. The RVD sequences of the dTALEs used in this study are provided in Table S5.

### 5’-RACE PCR analysis

Total RNA was isolated from leaves of Azucena 24 h after inoculation with *Xoo* strain MAI1 or *Xo* X11-5A (p*talB*) using the RNeasy kit (Qiagen), and subjected to 5’-RACE RT-PCR analysis using primer OsERF#123-Rev2 and the 5′ RACE System for Rapid Amplification of cDNA Ends (Invitrogen). Individual cDNAs were cloned in pGEM-T easy vector (Promega) and sequenced.

### Bioinformatics

Genomes of strain *Xoo* BAI3 and MAI1 were sequenced using SMRT whole genome sequencing (Icahn School of Medicine at Mount Sinai, New York, USA) and assembled using HGAP 3.0 as described [41]. Genomes were circularized using the AMOS package, specifically minimus2 under default parameters [64]. Genome sequences and raw sequences were deposited at the NCBI SRA (ncbi.nlm.nih.gov) under Bioproject PRJNA427174. TAL effector sequences were extracted from genomes using in-house perl scripts, and verified using the program PBX [43].

Genomes of other *Xanthomonas strains* were obtained from the NCBI under accession numbers: *Xoc* B8-12: CP011955.1, *Xoc* BLS279: CP011956.1, *Xoc* BLS256: CP003057.1, *Xoc* BXOR1: CP011957.1, *Xoc* CFBP2286: CP011962.1, *Xoc* CFBP7331: CP011958.1, *Xoc* CFBP7341: CP011959.1, *Xoc* CFBP7342: CP007221.1, *Xoc* L8: CP011960.1, *Xoc* MAI10: AYSY01, *Xoc* RS105: CP011961.1, *Xoo* CFBP1947: CP013666.1, *Xoo* KACC10331: AE013598.1, *Xoo* MAFF31101: AP008229.1, *Xoo* NAI8: AYSX01, *Xoo* PXO83 CP012947, *Xoo* PXO99^A^: CP000967.1, *Xoo* PXO86: CP007166.1, *Xoo* X11-5A: LHUJ01, *Xoo* X8-1A: AFHL01, *Xtu* CFBP2541: NZ_CM003052.1, NZ_CM003053.1.

Amino-acid sequences of housekeeping genes were obtained from each genome using AMPHORA [65], and then aligned using MUSCLE with default parameters v3.8.31 [66]. Poorly aligned positions were removed using Gblocks v. 0.91b [67]. Phylogenetic trees were then generated with PhyML v. 3.1 [68]. Trees were then visualized using the package ape from R [69]. ANI blast values were calculated for all pairs of genomes analyzed using the ANI calculator form the enveomics suite [44]. *In silico* DNA-DNA hybridization values were calculated using the Genome-to-Genome Distance Calculator version 1 [45]. Dot-plots of paired whole genome alignments were generated using Gepard v. 1.4 [70], with word-size = 30 and window= 30. Multiple genome alignments were made using progressive Mauve v. 2.4.0 [71]. Alignments were visualized using GenomeRing [72]. Circular genome representations were also made with BRIG v.0.95 [73]. Insertion Sequences families in *Xo* genomes were identified using IS-Saga [74]. In the case of Tn*Xax* 1-like sequences, nucleotide sequences were obtained from Ferreira *et al.* [46] and were aligned against *Xo* genomes using BLAST 2.2.28+ (blastn –evalue 0.0001 -word_size 7).

Sequences of non-TAL effectors were extracted from The Xanthomonas resource (http://xanthomonas.org) and aligned against *Xo* genomes using BLAST 2.2.28+ (blastn BLAST 2.2.28+). TAL effector sequences were extracted from genomes using in-house perl scripts. Repeat-based trees were generated using DisTAL with default parameters [47] and trees based on predicted DNA-binding specificity were created using FuncTAL with default parameters [47]. TAL effectors were also classified in families using the suite AnnoTALE with default parameters [36]. N-and C-terminal sequences of TAL effectors were aligned using MUSCLE with default parameters v3.8.31 [66] and phylogenetic trees were then generated with PhyML v. 3.1 [68] as above. Unique TAL effector central repeats were also identified using DisTAL [47], and repeat sequences were visualized in R using the heatmap.2 function. A vector of colors was created to represent each repeat using the rainbow function of R, based on distances between repeats calculated from Needleman-Wunsch nucleotide alignments obtained from DisTAL [47].

Target predictions were made for RVD sequences of African *Xoo* TAL effectors using Talvez v.3.1 [47] against the promoter region of annotated rice genes (Nipponbare v. MSU7), 1000bp before the translation start site. The top 600 genes with the highest EBE prediction score were kept for intersection with expression data to identify candidate targets for each TAL effector.

Microarray data used correspond to the Gene Expression Omnibus (GEO) dataset GSE19844. The Affymetrix probe-level expression data in CEL files was processed using the Bioconductor affy package [75] with default parameters of the rma function that computes the RMA (Robust Multichip Average) expression measure after background correction, normalization and probe summarization. The Bioconductor limma package was used to identify differentially expressed genes as described previously in [76]. The probesets that displayed a value of the log2-transformed fold change (logFC) equal or above 1 and an associated adjusted p-value (Benjamini and Hochberg’s adjustment method) equal or below 0.1 were considered as differentially expressed. The mapping of Affymetrix probe sets on MSU7 Gene Model IDs was obtained from the www.ricechip.org website.

Other routine computational tasks were performed in R with utilities provided by packages from the Bioconductor project [77]. All the expression data, RVD sequences for TAL effectors, and TAL binding sites predictions here included can be explored in our recently released tool daTALbase (http://bioinfo-web.mpl.ird.fr/cgi-bin2/datalbase/home.cgi) [78].

## Acknowledgments

This project was supported by grants from the Agence Nationale de la Recherche (http://www.agence-nationale-recherche.fr; ANR-14-CE19-443-0002toBS), the CGIAR Research Program on rice agri-food systems (http://www.grisp.net/main/summary; MENERGEP NewFrontier program to BS), the US National Science Foundation (https://www.nsf.gov; IOS-1238189 to AB). TTT and AP-Q were also supported by doctoral fellowships awarded by the Erasmus Mundus Action 2 PANACEA and PRECIOSA program of the European Community, respectively (http://eacea.ec.europa.eu/erasmus_mundus/index_en.php). We are grateful to J. M. Jacobs for valuable comments.

## Author Contribution

T.T.T., A.P.Q., M.H., S.C., I.W. and Y.Y. performed the experiments. T.T.T., A.P.Q., M.H, S.C., L.W., S.C., A.B., R.K. and B.S. analyzed the data. J.L. and V.V. contributed reagents/materials/analysis tools. T.T.T., A.P.Q., A.B., R.K., M.H. and B.S. planned and designed the research and wrote the manuscript.

## Supplementary Information

**Figure S1. Whole genome alignments of *X. oryzae* strains.** (A) Paired dot plots showing whole genome alignments between *Xo* genomes. Dotplots were made using Gepard [70] with word-size = 30 and window= 30. For visualization, the upper color limit was set to 50%, and the lower color limit and grayscale to zero. (B) Mauve multiple alignment of selected *Xo* genomes. (C) Heatmap shows average nucleotide identity (ANI) values for all pairs of genomes calculated using the ANI tool from the enveomics suite [44], top shows hierarchical clustering based on these values.

**Figure S2.Insertion sequences (IS) in *X. oryzae* genomes.** Frequency (total instances) of insertion elements identified in the genomes of African (MAI1, BAI3 and CFBP1947) and Asian (PXO99^A^) *Xoo* strains, and in *Xoc* strain BLS256. IS were identified using IS-Saga, and family names used in the IS-Saga database are shown [74]. Tn*Xax1*-like sequences were identified using BLAST.

**Figure S3.Shared predicted targets for groups of African *X. oryzae* pv. *oryzae* TAL effectors**. Heatmap showing the percentage of shared predicted targets for all pairs of African *Xoo* TAL effectors. Predictions were made using TALVEZ v.3.1 [62], and the top 500 predicted targets based on locus IDs were used for each comparison.

**Figure S4.*Os09g39810* and *Os09g29820* are induced by TalB_MAI1_**. sqRT-PCR analysis of *Os09g39810* (aka *OsERF#123*), *Os09g29820* (aka *OsTFX1*) and actin expression levels 24 hours post infiltration of Azucena rice leaves with *Xoo* strain MAI1, and *Xo*X11-5A strains carrying *talB*_MAI1_ or an empty vector (*EV*). RNA quality and quantity was estimated using a ND-1000 Nanodrop spectrophotometer. cDNA was synthesized with 1 μg of total RNA by means of the SuperScript first-strand synthesis system (Invitrogen, Carlsbad, CA, U.S.A.).

**Figure S5.*OsERF#123* transcription start sites observed in response to TalB**. RNA collected 24 hours post-infiltration of rice leaves with *Xoo* strains MAI1 and X11-5A (p*talB*) were subjected to 5’ RACE. Obtained cDNA sequences were aligned above (MAI1) and below (X11-5A (p*talB*)) the predicted 5’-UTR of *OsERF#123*. The effector binding element (EBE) for TalB is highlighted in gray. Numbering starts downstream of the 3’ end of the EBE. The total number of sequences obtained (N) is given on the left.

Table S1.African *X. oryzae* pv. *oryzae* TAL effector nomenclature.

Table S2.Candidate targets for MAI1 TAL effectors in the rice genome.

Table S3.Candidate targets for BAI3 TAL effectors in the rice genome.

Table S4.Primer sequences used in this study.

Table S5.RVD sequences of the dTALEs used in this study and their target sequence.

